# VelosGT: Edge-Computing for Human Variant Analysis

**DOI:** 10.1101/2023.08.11.550418

**Authors:** Josh Flygare, Alex Born

## Abstract

Velos^GT^ is a MacOS application to analyze human genomes for genetic mutation and rare disease. It consumes raw read files (fasta/fastq) and returns annotated, functionally prioritized variants in a single step, while significantly out-performing standard pipelines. Velos^GT^ ingests short-read data in all common formats (fastq, fasta, fastq.gz, and fasta.gz) as well as paired data. It can perform analyses on single genomes (proband) or trios. The underlying algorithms have been optimized to the extent that it can run directly on a personal computer, with no additional hardware required. This increases convenience and speed because no large uploads or downloads are required. All discovered variants are immediately displayed and searchable within the app. On an average^*^ Mac laptop, Velos^GT^ will return the results of a high-coverage 30GB read file in under an hour, and a 5GB read file in ten minutes or less. Trios can be computed using the completed child-parent analyses in a few minutes.

## Introduction

Healthcare professionals and researchers need useful, relevant results. The VelosGT pipeline can recover both novel and clinically known variants within all known genes in the human genome. Once the analysis is complete, the variants are presented in a sortable, searchable table. Users can search for variants using many attributes including position, gene name, amino acid, pathogenicity, and even disease name, symptoms, and phenotype. As an example, if a neurological disorder or lymphoma is suspected, a search of ‘neuro’ or ‘lymph’ will return all sample variants associated with those keywords.

A direct performance comparison of a standard pipeline composed of the BWA aligner, samtools pileup, and freebayes variant caller is shown in Figure 1. Velos^GT^ consistently out-performs the standard pipeline by about 100%. Performance gains are especially prominent for high-coverage files containing many reads.

**Figure 1:**
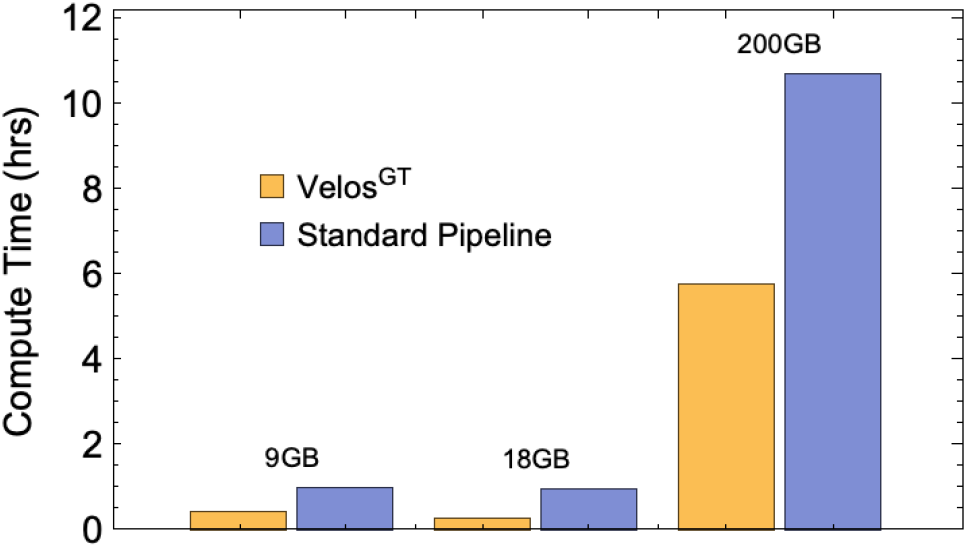
On average, Velos^GT^ outperforms an example genetics pipeline implementing BWA/samtools/freebayes by about 100%. Alignment and variant calling of BWA and freebayes were parallelized for a more direct comparison. File size (given in uncompressed GB) is less important to overall compute time as compared to number of reads aligned, as seen in the similar time of the first two datasets (9GB and 18GB).

The accuracy of VelosGT was validated using read simulations containing variants from ClinVar. VelosGT detects SNPs, MNPs, indels and clumped variants. All variants 50bp or greater are considered structural variants (SVs) and are excluded from the simulation. In total, the analysis pipeline covers about 95.3% (1.4 million) of the chromosomal variants in ClinVar. The tests included SNPs, MNPs, indels, and clumped variants, of which Velos^GT^ recovered more than 99.3%. The Verification and Performance section discusses these simulations in depth. Case studies were then used to show the app’s sensitivity and successful recall of functionally relevant variants.

### Finding the Variants That Matter

Velos^GT^ discovers functionally-relevant variants occurring within all known genes by comparing reads against the GRCh38 human genome reference. About 85% of known pathogenic variants are contained within the human exome [2]. The other 15% are spread through intronic regions of the genome. These regions contain vital gene activity regulators, changes to which can cause genes to be silenced or transcribed unexpectedly. Velos^GT^ is optimized to search these gene regions. These resulting variants are annotated with metadata which can be used to determine the functional effects.

In every individual’s genome there are millions of variants, of which many are novel, having yet to be attributed to a functional change. Sifting through millions of possibly innocuous changes to find the known pathogenic variants requires sophisticated tooling [1]. Velos^GT^ contains more than two billion annotations for SNPs and indels to identify implicated variants. The annotations include the name of the gene that the variant affects, the amino acid change which is induced, as well as a number of other attributes (see Appendix for complete list). The annotations can be used to determine whether the variant is in an intergenic region, is part of the transcriptome, is coding, or affects a regulatory feature. Variants within coding regions are assigned SIFT and PolyPhen scores which predict how an amino acid substitution affects protein expression. The SIFT scores predict these changes on the basis of sequence homology and physico-chemical similarly. PolyPhen scores predict the impact of these substitutions on the basis of phylogenetic and structural information. These scores can be used to estimate the likelihood a that change will be benign or deleterious.

Additionally, Velos^GT^ searches for variants in ClinVar, a database which aggregates information about how genetic variants impact health outcomes. Velos^GT^ can match against both normalized and non-normalized ClinVar variants. If a matching variant is found, twenty additional data fields including disease names, pathogenicity and possible symptoms are attached to the variant. (See the appendix for more details.) Velos^GT^ is also able to detect and functionally annotate overlapping indels. Velos^GT^ combines annotations, integration with ClinVar, and call-level information like indel overlap in a single convenient app, giving clinicians the necessary tools to find the variants that matter most.

### Discovering Inheritance (trio analysis)

After files have been processed for variants, a user can select 3 completed samples to use in a trio analysis. The app computes the Mendelian inheritance of all variants in the samples, including autosomal and sex chromosomes. The inheritance of the resulting variants are returned with likelihood estimates computed using a Bayesian model based on variant zygosities. Trio variants, depending on the phenotype of the parents and child, can be categorized as dominant, recessive, indeterminate, or as a violation. Dominant and recessive are computed using the standard definitions. Indeterminate means it is not possible to differentiate between dominant and recessive given the selected phenotypes and computed zygosities. A ‘violation’ means the combination of zygosities, phenotypes, and chromosomal location resulted in a violation of the principles of Mendelian inheritance. Likely, in practice, this is due to regions with very poor coverage or homologous regions that result in mapping difficulties.

All trio variants are displayed within the app tables for sorting and searching. After processing the initial trio, the user can change phenotypes of the male parent, female parent, or child to investigate other diseases. The disease finder tool also works with trio variants to help narrow the results to those that are most functionally relevant.

### Computational Performance

The performance of Velos^GT^ was assessed by running samples on different machines. Estimates for the time to perform an analysis (wall clock time) are shown in Table 1. The file size is a good estimator of the total processing time and is a measurement (in bytes) of the uncompressed read data. The CPU is the primary metric in the computation time. During internal testing, memory usage never exceeded 4 GB. On modern computers, users can continue to use their computer for normal tasks while Velos^GT^ performs analyses.

**Table 1:** Computed and estimated processing times in minutes for various size files, where sizes are uncompressed gigabytes.

| CPU | Processing Time (min) |  |  |  |
| --- | --- | --- | --- | --- |
|  | 0.85 GB | 2.2 GB | 30 GB | 150 GB |
| M1 Pro | 0.37 | 1.5 | 12.3 | 87(1.5hr) |
| M1 | 0.46 | 1.86 | 15.4 | 108(1.8hr) |
| 2019 I7 | 0.68 | 2.27 | 17.6 | 117(2.0hr) |
| 2015 I7 | 0.96 | 8.27 | 31 | 206(3.4hr) |

### Velos^GT^: The Computational Tool

The Velos^GT^ app performs three major steps in processing read files, 1) pre-processing and alignment, 2) variant calling and prioritization (including association with ClinVar variants), and 3) annotation of all possible SNVs and many indels. These steps constitute a bioinformatics strategy and pipeline that achieves excellent throughput even on limited hardware, such as a laptop or Mac mini.

#### Variant Calling and Prioritization

The variant caller for Velos^GT^ is built to be highly sensitive and reliable, returning variants whose allele frequency exceeds a minimum threshold. We determined this threshold using statistical techniques to optimize between sensitivity and specificity. The app lists a calling quality for each variant. A call quality of ‘high’ represents a precision greater than 0.97, whereas ‘nominal’ represents a precision of 0.88. These thresholds were determined from a 30x Illumina read sample from the Genome in a Bottle HG002 benchmark. When calling a variant, there are many factors that affect confidence such as read quality, mapping quality, soft clipping, etc. While appropriately accounting for these parameters can be helpful to increase specificity, over-reliance on them can lead to a decrease in sensitivity, potentially missing key variants.

We chose to tune Velos^GT^ to be highly-sensitive and predictable because 1) Velos^GT^ focuses the analysis on known genes, and 2) the user can choose how to narrow the search space using the tools provided in the app. This decreases the possibility of missing a variant and in research, is critical to avoid. The app also displays variants for which there is sub-optimal evidence in the sample reads, and these variants should be investigated further. After discovering a variant using the app, it is recommended that the results be confirmed using independent analyses or assessments.

Calling SNVs and indels is relatively straight-forward compared to more complex variants, such as insertion-deletion combinations. There are nearly 2000 variants in ClinVar represented by these composite insertion-deletions. Depending on the mapping choice of the aligner, the variants called can appear different from the ones presented in literature or variant databases. Velos^GT^ resolves common ambiguities in non-normalized representations to match variants against the representation found in variant databases.

#### Security Considerations

Velos^GT^ stores the analysis results in the in-app library. This library resides in the application support directory on the user’s computer. Enabling MacOS’s build-in FileVault disk encryption can protect these analyses and source read data from unauthorized access. This stops individuals with physical access (e.g. a computer thief or a rouge employee) from accessing data on the hard drive. In addition to enabling FileVault, users should follow standard security procedures including the use of strong passwords, the application of security updates, and the creation of strong network security policies.

#### Verification and Performance

To test the reliability and sensitivity of the variant caller, as well as the performance of the internal pipeline, we simulated the chromosomal variants found in ClinVar. Our simulation included SNPs, indels, MNPs, and clumped variants up to 50 base pairs in length that are within the regions of interest. In total, Velos^GT^ covers about 1.4 million of the 1.45 million chromosomal variants in ClinVar, or 95.3%. Initially, to estimate the ideal recovery rate for non-trival variants such as clumped variants or overlapping indels, we inserted all 130,000 indels and clumped variants into GRCh38. This was done with selective batches to avoid variant collisions. The recovery rate of indels and clumped variants (up to 50 bp) were 99.78% and 91.45%, respectively. These numbers can be viewed as ‘ideal condition’ recovery rates.

To estimate the recovery rate for SNPs, indels, MNPs, and clumped variants in non-ideal conditions, we used the Mason Simulation package to produce read pileups with industry standard error rates for Illumina 70bp, 150bp and 200bp reads mason. The error rate curves can be found in the previous reference, but typical values for positional mismatch errors are about 1% while insertion and deletion rates are closer to 0.04%. Over 40,000 variants were randomly chosen from ClinVar, spread across the entire genome. Velos^GT^ was then used to recall these variants, with the results shown in Table 2.

**Table 2:** ClinVar recovery using 200bp (top), 150bp (middle), and 70bp (bottom) Illumina paired-end reads with industry standard positional and indel error rates.

| Recovery Rate |  |  |  |
| --- | --- | --- | --- |
| Type | n Simulated | Missed | % Recovered |
| 200bp |  |  |  |
| SNP | 36235 | 86 | 99.76 |
| indel | 5232 | 68 | 98.70 |
| MNP | 211 | 1 | 99.53 |
| Clumped | 259 | 34 | 86.87 |
| 150bp |  |  |  |
| SNP | 36235 | 96 | 99.74 |
| indel | 5232 | 125 | 97.61 |
| MNP | 211 | 1 | 99.53 |
| Clumped | 259 | 34 | 86.87 |
| 70bp |  |  |  |
| SNP | 36235 | 50 | 99.86 |
| indel | 5232 | 459 | 91.23 |
| MNP | 211 | 2 | 99.05 |
| Clumped | 259 | 52 | 79.92 |

To understand the effect read length has on predicted recovery rates, we then simulated that same random subset of variants with 200bp 150bp and 70bp Illumina paired-end reads. As discussed previously, longer reads are more mappable, and less susceptible to damage from longer variants. We would expect the recovery rates for 200bp reads to be slightly better than 70bp or 150bp. This turns out to be true on average, as seen in Table 2.

The primary source of error in these simulations are issues with mapping reads in highly homologous regions. It is expected that the 200bp reads also perform much better with larger indels and clumped variants, as the reads are more capable of surviving a 50bp change while still being mappable. The histograms in Figure 3 show the lengths of recovered variants (indels and clumps) for the 70bp and 200bp reads. The 200bp reads, as expected, recovered larger indels and clumped variants, up to 50bp. The 70bp reads were unable to recover variants larger than 30bp, but performed within a few percent of the 200bp reads under about 20bp variant length.

**Figure 2:**
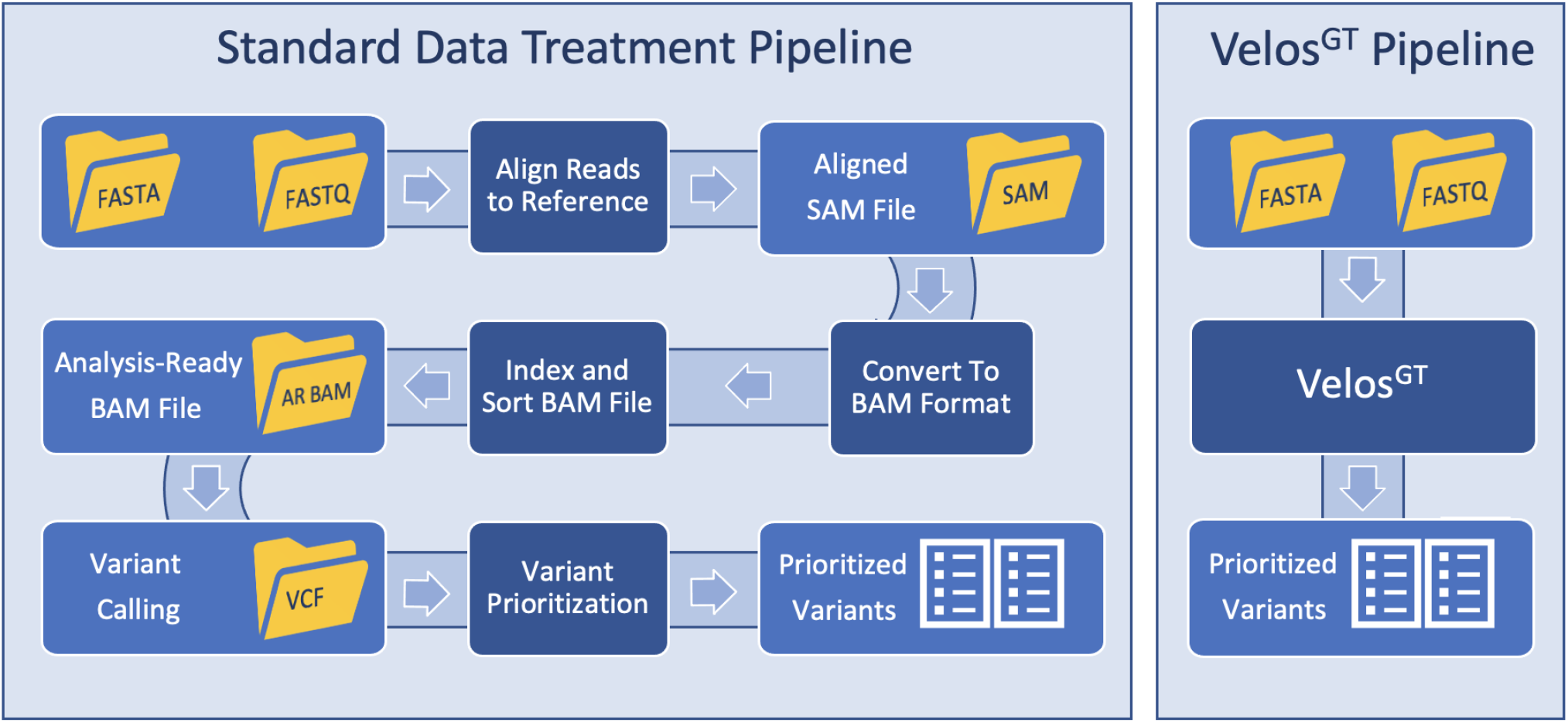
A comparison of a standard variant analysis pipeline (left) to the Velos^GT^ pipeline (right).

**Figure 3:**
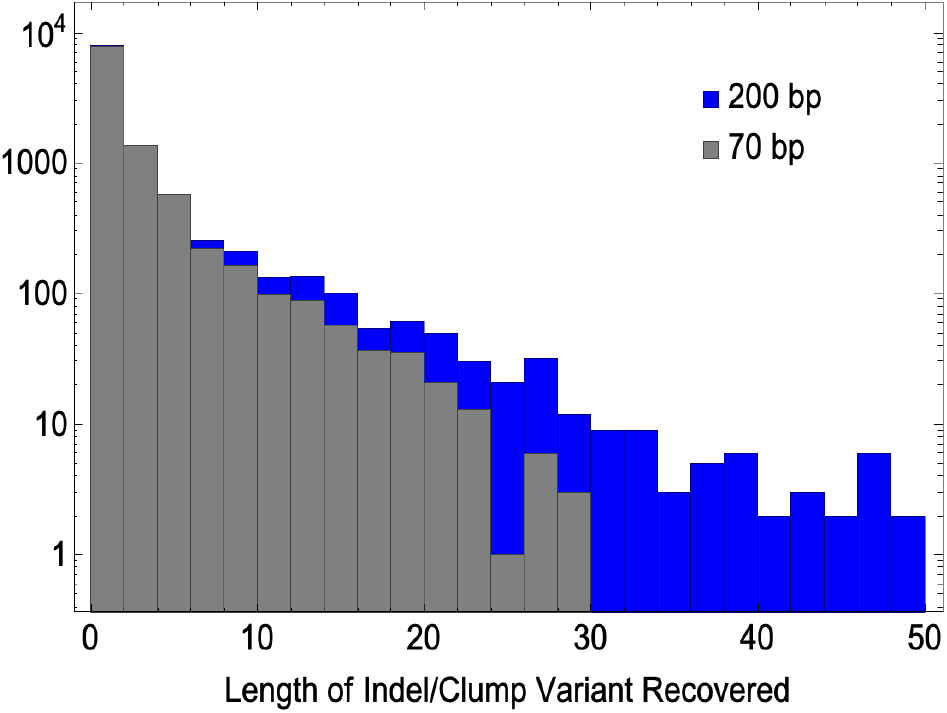
Histograms of recovery rate of large variants, comparing 200bp (blue) to 70bp (gray).

#### Quick Examples

To illustrate use cases of the app and some of its features, the results of two publicly available samples are discussed below. The NCBI SRA ID is listed for each, and small samples were chosen so users can easily replicate the results if desired.

The first clinical sample was known to have Tay-Sachs disease (from NCBI SRA SRR15180921). In this particular sample, being a relatively small gene panel subset with very deep coverage, Velos^GT^ returned two variants, both of which were found in ClinVar, but only one of which was pathogenic. The app table and variant detail viewer showed it was an insertion of GATA in chromosome 15 within the HEXA gene at a variant allele frequency of 0.92. The annotation confirmed it as a frameshift mutation with a consequence score of seven.

The next sample was a known case of dilated cardiomyopathy (NCBI SRA SRR19964385), which is typically caused by a mutation in the TTN (Titin) gene. The compressed read file of the exome was 127 MB, and returned about 5900 variants. Using the app’s ‘disease finder’ feature, which lists all diseases from ClinVar associated with variants in the sample, ‘Dilated cardiomyopathy 1G’ had 52 associated variants. Using the advanced search tool with ‘cardiomyopathy’ for diseases/symptoms and ‘ttn’ for gene, only one variant remained. It’s clinical significance was listed as ‘conflicting interpretations of pathogenicity’. The allele frequency for the remaining variant was 0.48, with a high probability of being heterozygous.

## Appendix

| Annotation Details |  |
| --- | --- |
| Type | SNV, INS, DEL |
| Length | Length of variation (SNV = 0, an INS like GA = 2) |
| Annotation Type | Regulatory Feature, Transcript, Coding Transcript, Intergenic, Non-Coding Transcript |
| Consequence | Regulatory, Intronic, Splice Site, Canonical Splice, Non-Synonymous, Stop Gained, Synonymous, Downstream, Non-coding Change, Upstream, Stop Lost, 3-prime UTR, 5-prime UTR, Intergenic |
| Consequence Score | Range 0.0 - 8.0, higher number is more likely to be deleterious. |
| Consequence Details | Description of all consequential features of variant (e.g. a single variant may return 'intron, 5-prime UTR, splice acceptor, coding sequence') |
| Original Amino Acid | Uses international amino acid abbreviations (see https://www.ddbj.nig.ac.jp/ddbj/code-e.html#amino-1) |
| New Amino Acid | Same format as Original Amino Acids |
| Gene ID | The Gene ID of the genes containing the variant, as referenced on sites like https://www.genecards.org/ |
| Feature ID | The ID of the sequence feature containing the variant. |
| Gene Name | Can also be referenced on gene cards. |
| CCDS | Consensus CDS ID, as can be found at https://www.ncbi.nlm.nih.gov/projects/CCDS/CcdsBrowse.cgi |
| SIFT Category | Categories for the SIFT score. Tolerated, deleterious. |
| SIFT Score | Range 0.0 - 1.0, where 0.0 is most likely to be damaging, and 1.0 is predicted to be benign. |
| PolyPhen Category | Categories for Polymorphism Phenotyping Score. Benign, probably damaging, possibly damaging, unknown. |
| PolyPhen Score | Range 0.0 - 1.0, where 0.0 is most likely to be benign, and 1.0 is predicted to be damaging. |

| ClinVar Details |  |
| --- | --- |
| Allele Frequencies (ESP) | Allele frequencies from GO-ESP https://evs.gs.washington.edu/EVS/ |
| Allele Frequencies (ExAC) | Allele frequencies from ExAC https://gnomad.broadinstitute.org/ |
| Allele Frequencies (TGP) | Allele frequencies from TGP https://www.ncbi.nlm.nih.gov/projects/gap/cgi-bin/study.cgi?study_id=phs000495.v1.p1 |
| Allele ID | ClinVar Allele ID, can be searched at https://www.ncbi.nlm.nih.gov/clinvar/ |
| Disease Name | ClinVar's preferred disease name for the concept specified in the disease identifiers |
| Disease Name (FIV) | ClinVar's preferred disease name for included variant (FIV) specified in disease identifier FIV |
| Disease Identifier | Tag-value pairs of disease database name and identifier, e.g. OMIM:NNNNN |
| Disease Identifier (FIV) | Tag-value pairs of disease database name and identifier for included variant, e.g. OMIM:NNNNN |
| Top-Level HGVS expression | Human genome variation society standard format https://www.ncbi.nlm.nih.gov/clinvar/docs/hgvs_types/ |
| Review Status | ClinVar review status for the Variant ID. |
| Clinical Significance | The clinical significance for the individual variant. Multiple values are separate by vertical bar |
| Conflicting Clinical Significance | Also for individual variant, multiple values separated by vertical bar |
| Haplotype/Genotype Clinical Significance | Reported as pairs of VariationID:clinical significance, also separates multiple values with vertical bar |
| Variant Type | SNV, deletion, insertion, duplication, microsatellite, variation, inversion, copy number loss, copy number gain |
| Sequence Ontology ID | IDs can be searched at http://www.sequenceontology.org/browser/obob.cgi |
| Clinical Sources | Represented as tag value pairs of database:variant identifier. |
| NSV Accessions from db-Var | Can be found at https://www.ncbi.nlm.nih.gov/dbvar/ |
| Gene Info | The gene symbol and ID separated as a tag value pair. These are searchable in the app and automatically link to GeneCards. |
| Molecular Consequence | List of molecular consequence as tag value pair of Sequence Ontology ID and molecular consequences, separated by vertical bar |
| Allele Origin | Multiple types including germline, somatic, unknown, inherited, parental, maternal, de-novo, biparental, uniparental, not-tested, testing inconclusive, other. |
| RS Number | dbSNP ID, can be searched at https://www.ncbi.nlm.nih.gov/projects/SNP/snp_summary.cgi |

